# 3D bioprinting of an implantable xeno-free vascularized human skin graft

**DOI:** 10.1101/2022.02.21.481363

**Authors:** Tania Baltazar, Bo Jiang, Alejandra Moncayo, Jonathan Merola, Mohammad Z. Albanna, W. Mark Saltzman, Jordan S. Pober

## Abstract

Bioengineered tissues or organs produced using matrix proteins or components derived from xenogeneic sources pose risks of allergic responses, immune rejection, or even autoimmunity. Here, we report successful xeno-free isolation, expansion, and cryopreservation of human endothelial cells, fibroblasts, pericytes and keratinocytes from a single donor. We further demonstrate the bioprinting of a human skin substitute with a dermal layer containing xeno-free cultured human endothelial cells (EC), fibroblasts, and pericytes in a xeno-free bioink containing human collagen type I and fibronectin layered in a biocompatible polyglycolic acid (PGA) mesh and subsequently seeded with xeno-free human keratinocytes to form an epidermal layer. Following implantation of such bilayered skin grafts on the dorsum of immunodeficient mice, keratinocytes form a mature stratified epidermis with rete ridge-like structures. The ECs and pericytes form human EC-lined perfused microvessels within 2 weeks after implantation, preventing graft necrosis, and eliciting further perfusion of the graft by angiogenic host microvessels. In summary, we describe the fabrication of a bioprinted vascularized bilayered skin substitute under completely xeno-free culture conditions demonstrating feasibility of a xeno-free approach to complex tissue engineering.

## INTRODUCTION

Bioengineered skin substitutes are approved for clinical use in patients with non-healing skin ulcers or surgically resected skin cancers, but all currently available products contain xenogeneic components, either in the form of matrix molecules or human cells that have been expanded in culture using media supplemented with xenogeneic components such as fetal bovine serum (FBS)^1–3^. Despite promising clinical outcomes, some patients have exhibited allergic responses. For example, Apligraf™, a bilayered skin graft manufactured with allogeneic keratinocytes and dermal fibroblasts along with bovine collagen, is contraindicated in patients with known allergies to bovine collagen^4,5^. Moreover, patients grafted with cultured human keratinocytes, for burn wounds, have been shown to make antibodies to xenogeneic antigens such as FBS, which presumably derived from culture medium^6,7^. Antigen persistence can potentially make such human keratinocytes susceptible to immune destruction by recognition of proteins in FBS and thereby contribute to delayed loss of human keratinocyte grafts. Previous efforts have been made to reduce keratinocyte immunogenicity by growing cells in a serum-free medium with bovine pituitary extract as a serum substitute^8^. However, bovine pituitary extract contains non-human proteins so that human keratinocytes cultured in medium that is supplemented with bovine pituitary extract could similarly activate an immune response. In addition, species specific variation between human and animal sources of collagen type I can substantially alter the physicochemical properties of collagen, which in turn can induce a cellular immune response^9,10^. Finally, xenogeneic proteins in serum or matrix can break tolerance to human homologous proteins, leading to autoimmunity^11–13^. Consequently, health authorities currently recommend the use of ‘xeno-free’ protocols in the clinical translation of skin equivalents^14^. However, to our knowledge, generation of a fully xeno-free complex tissue capable of surviving after implantation into a host has yet to be demonstrated.

Another challenge in achieving successful clinic translation of bioengineered tissues and organ is vascularization^15^. Notably, Apligraf and other approved bilayered skin replacements fail to permanently engraft because they lack dermal vascular networks important for perfusion^16,17^. In previous experiments using skin constructs generated by repopulation of decellularized dermis we showed that allowing human ECs to infiltrate the dermis led to microvessel formation and early perfusion following engraftment on the dorsum of immunodeficient mice, thereby improving graft survival^18^. More recently we demonstrated that inclusion of human vascular cells into the dermal bioink used for 3D printing of skin also produced a bi-layered, vascularized skin construct that is perfused, through both graft and host microvessels, early after implantation on immunodeficient mice^19^. Building upon this work, we now demonstrate the fabrication of a bioprinted vascularized skin graft generated under strict xenogeneic-free conditions starting from isolation, culture and cryopreservation of primary cells, bioink/scaffold development, construct maturation *in vitro*, and successful engraftment on an immunodeficient mouse host. We also improve the mechanical properties of our previously described 3D printed graft by incorporation of a biodegradable PGA mesh into the dermal layer.

## MATERIALS AND METHODS

### 1.1 Cell isolation, expansion, and cryopreservation of primary human cells under xeno-free conditions

Endothelial cell (EC) cultures were established from human endothelial colony forming cells (HECFCs) present in human umbilical cord blood by culturing whole cord blood mononuclear cells from de-identified donors in EC-Cult-XF medium (STEMCELL Technologies) supplemented with 2 mg/mL heparin (#07980; STEMCELL Technologies) on 10 μg/mL human plasma fibronectin (#FC010, EMD Millipore)-coated plates, as previously described. EC colonies spontaneously arose as “late outgrowth cells” by day 14, were allowed to reach confluence, and were then serially passaged in the same medium. Human microvascular placental pericytes (PCs) were isolated from discarded and de-identified placentas as explant-outgrowth cells in EC-Cult-XF supplemented with 2 mg/mL heparin on 10 μg/ml human plasma fibronectin-coated plates, also as previously described ^20^. Human fibroblasts (FBs) and keratinocytes (KCs) were isolated from freshly discarded human foreskin samples according to previously published protocols ^19^. FBs obtained from the dermis and KCs obtained from the epidermis were cultured in FibroLife Xeno-free Complete medium (#LM-0013; LifeLine Cell Technologies) and EpiLife medium (#MEPI500CA; ThermoFisher) supplemented with HKSdaFREE and HKGE (AvantBio), respectively. KCs were cultured and serially passaged into 7 μg/cm^2^ human collagen IV-coated plates (#C7521; Sigma-Aldrich). All cell sources were obtained at Yale-New Haven Hospital under protocols approved by the Yale University Institutional Review Board including, where indicated, all four cell populations were obtained from a single donor source through the Yale Department of Obstetrics and Gynecology. All primary cells were used between subculture 3–7 and were routinely cryopreserved in 90% human serum (#H4522; Sigma-Aldrich) supplemented with 10% dimethyl sulfoxide (DMSO).

### 1.2 Trans-endothelial electrical resistance (TEER) measurements

EC proliferation and barrier formation were assessed by measurements of trans-endothelial electrical resistance (TEER) in an electrical cell-substrate impedance system (ECIS), a technique for continuous measurement of barrier integrity^21^. Resistances were measured by application of a 1 μA constant alternating current (AC) at 4000 Hz between a large and small electrode embedded in the chamber slide and recorded by an ECIS Z-theta instrument (Applied BioPhysics). Briefly, 1 x 10^4^/mL ECs were plated on 8-well gold electrode arrays (catalog # 8W10E+ PET; Applied BioPhysics). Culture medium, EGM2-MV and EC-Cult-XF, was changed every other day, and TEER measurements were obtained every 10 hours. To assess cytokine-induced changes in TEER, 1 ng/mL human recombinant tumor necrosis factor-α (TNF; PeproTech) was introduced to the medium after confluency was achieved and the TEER was recorded every 2 hours. Raw data were exported from ECIS software as .CSV extension files and imported into Prism software for plotting and analysis.

### 1.3 Phenotyping of primary cell cultures

ECs, FBs, PCs and KCs cultured under xeno-free conditions were analyzed for surface and intracellular markers expression by flow cytometry and immunofluorescence microscopy using antibodies against: CD31 (WM59; Biolegend), CD45 (2D1; Biolegend), ZO-1 (sc-33725; Santa Cruz), VE-cadherin (sc-6458, Santa Cruz), vWF (ab201336, abcam) claudin-5 (34-1600, Invitrogen), PDGFR-α (16A1; Biolegend), PDGFR-ß (18A2; Biolegend), CD90 (5E10, Biolegend), NG2 (9.2.27, Invitrogen), a-SMA (1A4, Invitrogen), FAP (AF3715, Novus Biologics), integrins α2β1 (ab24697, abcam), α5β1 (NBP2-52680; Novus Biologics), αvβ3 (sc-7312; Santa Cruz), α6 (MAB13501; R&D systems), α3 (MAB1345; R&D systems), β4 (NBP1-43369), CK14 (LL002; Novus Biologics), CK10 (EP1607IHCY, abcam) and occludin.

### 1.4 Preparation of xeno-free dermal and epidermal bioinks

Xeno-free dermal and epidermal bioinks were modified from a previously described protocol, substituting human for xenogeneic matrix proteins^19^. Specifically, the xeno-free dermal bioink was formulated with 4.5 x 10^6^ /mL human FBs, 4.5 x 10^6^ /mL human ECs and 5.5 x 10^5^ /mL human PCs, suspended in a solution prepared in the following order: 30 μL of human male AB serum (Sigma-Aldrich), 60 μL human plasma fibronectin (EMD Millipore), 60 μL of 10X HAM-F12 medium (Gibco), 330 uL of 6 mg/mL of human collagen type I (Humabiologics, Inc), 60 μL of 10x pH reconstitution buffer (0.05 M NaOH, 2.2% NaHCO3, and 200 mM HEPES) and 60 uL VitroGel Hydrogel Matrix (The Well Bioscience). VitroGel, a xeno-free hydrogel, was added to the bioink to minimize *in vitro* gel contraction and accelerate gelation of the dermal bioink. The ratio of VitroGel to human collagen type I in the dermal bioink was optimized *in vitro*. The condition that allowed for faster gelation while promoting self-assembly of EC-networks was selected (data not shown). The xeno-free epidermal bioink was formulated with 2 x 10^6^/ mL human KCs in 500 μL of EpiLife medium (#MEPI500CA; ThermoFisher) supplemented with HKSdaFREE and HKGE (AvantBio) and 5 ng/mL human recombinant animal-free KGF (Sigma-Aldrich), 1.64 mM CaCl_2_ and 50 ug/mL ascorbic acid. All cells were maintained at 37°C in a 5% CO_2_ atmosphere until printing. Where indicated, cultured cells were compared to their counterparts isolated and cultured using previously conditions involving use of non-human-derived materials^19^.

### 1.5 3D bioprinting of xeno-free skin equivalents

Skin grafts were fabricated using a commercially available bioprinter (model BioX; CELLINK). An exchangeable print head comprising a sterile 25-gauge stainless steel blunt needle (CELLINK) and maintained at 4°C was used to print the dermal bioink at extrusion pressure of 50 kPa. Skin constructs were generated by dispensing 0.2 mL dermal bioink containing FBs, PCs, and ECs, dispensed from a 3 mL syringe into a 24-well plate (Figure 1). A sterile polyglycolic acid (PGA) mesh (Confluent Medical Technologies, Inc.; thickness: 1 mm; density 50-60mg/cm, ᴓ: 13 mm) was added on top of the first printed layer and incubated at 37°C. The mesh was sterilized prior to printing by immersion in a 70% ethanol solution for 30 minutes and allowed to dry for 1 hour at RT. Upon gelation, a second layer of 400 μL of the dermal bioink was printed on top of the PGA mesh, incubated at 37°C until complete gelation, and submerged in EC-Cult XF media until the next day. Constructs (approximately 2 cm^2^ in size) were transferred to 6-well plates and submerged in a larger volume of media (4 mL) to prevent rapid exhaustion of nutrients. Medium was changed daily. After 4 days under EC-Cult-XF medium submersion to allow self-assembly of endothelial networks, the dermal construct was carefully transferred to a 24-well plate and the epidermal bioink containing KCs (1 x 10^6^ in 500 μL) was printed from a 3mL syringe, 30-gauge stainless steel blunt (CELLINK), at a pneumatic pressure of 20 kPa, on top of the dermal construct. Twenty-four hours after incubation at 37°C, the skin equivalents were transferred to a 6-well plate with EpiLife medium supplemented with HKSdaFREE, HKGE, 5 ng/mL KGF, 1.64 mM CaCl_2_, and 50 ug/mL ascorbic acid. The medium was changed every other day for 3 additional days.

**Figure 1.**
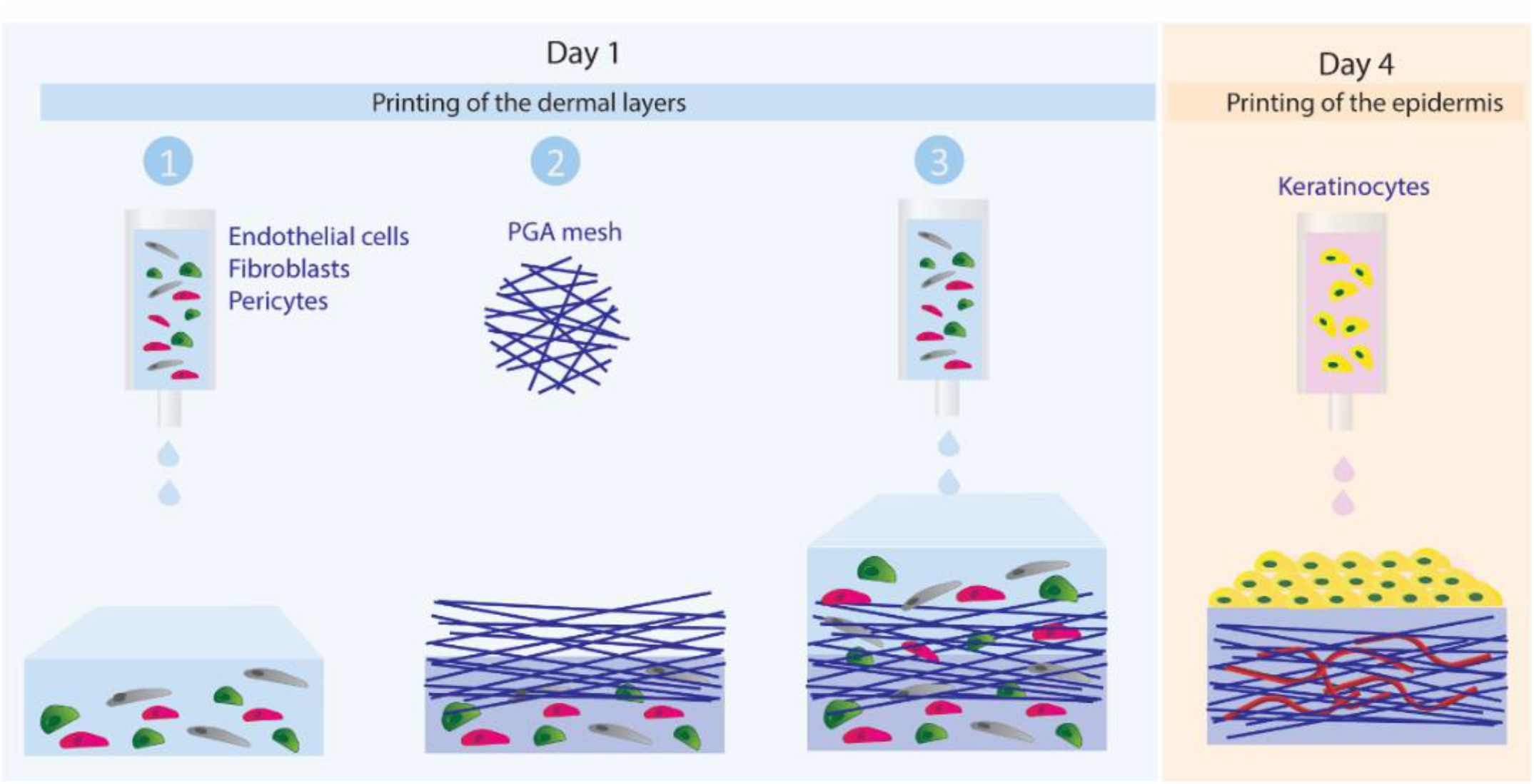
Schematic showing layer-by-layer bioprinting of xeno-free human skin equivalents.

### 1.6 Engraftment of bioprinted vascularized skin in SCID/bg mice

All procedures were performed under protocols approved by the Yale Institutional Animal Care and Use Committee. Twenty-seven printed skin constructs were engrafted under sterile conditions onto the side of twenty-seven 6-12 week old female C.B-17 SCID/bg mice (Taconic Farms, Germantown, NY) anesthetized by intraperitoneal injection of ketamine/xylazine. Mouse skin (approximately 1cm diameter) was excised from the dorsal side of the animal and a comparable sized piece of printed skin was placed on the wound and sutured in place using 6-0 Prolene suture. The graft was then covered with two layers of Vaseline gauze, pre-coated with Bacitracin cream, a layer of Tegaderm, two bandages covering the size of the wound and finally wrapped with Coban 3M tape. Bandaging was removed 10 days after implantation.

### 1.7 Assessment of *in vivo* perfusion by tail vein injection of fluorescein-UEA-I

Two-hundred microliters of 1 mg/mL fluorescein-conjugated Ulex europaeus agglutinin-I (UEA I; Vector) solution were injected per mouse via the tail vein and allowed to circulate for 30 min before harvesting grafts. Mice were euthanized and grafts were harvested to include surrounding and underlying mouse tissue and bisected perpendicular to the skin surface: one half of each graft was fixed in 10% buffered formalin overnight for paraffin-embedding; the other half was embedded in OCT, snap frozen, and cryo-sectioned (10 μm thick).

### 1.8 Morphological characterization of bioprinted skin grafts

Frozen sections (10 μm) of printed grafts were analyzed by H&E, immunofluorescence, and immunohistochemical staining. For immunofluorescence analysis, tissue sections were immersed in cold acetone for 10 minutes and blocked for 1 h with 10% normal goat serum in PBS or 10% donkey serum. Slides were then incubated with antibodies to cytokeratin 10 (rabbit, 1:200, clone EP1607IHCY; Abcam), cytokeratin 14 (mouse, 1:200, clone LL002; Abcam), collagen IV (mouse, 1:200, clone COL-94; Abcam), laminin 5 (rabbit, 1:100, Abcam), ki67 (rabbit, 1:200, clone SP6, Abcam), human CD31 (mouse, 1:200, clone JC70A, Agilent), mouse CD31 (rat, 1:200, clone SZ31, Dianova), mouse F4/80 (rat, 1:50, clone BM8, eBioscienceTM), or involucrin (mouse, 1:50, clone SY5, Abcam). Slides were then washed with PBS (3x) and incubated for 1h at room temperature with anti-mouse or anti-rabbit IgG H&L secondary antibodies conjugated with Alexa Fluor™ 488 or Alexa Fluor™ 568 (goat, 1:500; Abcam). Slides were mounted with VECTASHIELD antifade mounting medium containing DAPI (Vector Labs) for nuclear staining and imaged on a Leica DMI6000 inverted fluorescence microscope. Immunohistochemical analysis was performed with the following Biotin-SP AffiniPure IgG (H+L) antibodies: donkey anti-rabbit IgG (H+L), donkey anti-mouse IgG (H+L), and goat anti-rat. Substrate to the secondary antibody was included in the Vectastain ABC Peroxidase kit and AEC Peroxidase Substrate Kit (Vector Laboratories). In some cases slides were incubated with fluorescein-conjugated Ulex Europaeus Agglutinin I (UEA-1, 1:200, Vector Laboratories) to identify human ECs not labeled by perfusion.

### 1.9 Quantification of perfused human vessels, murine vessels, and non-perfused human vessels

Microscope images of stained tissue sections were taken using a Leica DMI6000 inverted fluorescence microscope. Five to ten randomly selected sections sampling all parts of the tissue block were evaluated per animal. Whole sections were imaged using a 10× objective, with an image pixel size of 0.916 μm. ImageJ software was used to quantify infused fluorescein ulex+, human CD31+ or mouse CD31+ area in each ×10 field section. Briefly, each corresponding channel image was binarized by defining a threshold equally applied to all images. The human perfused vessel area was defined as the number of infused fluorescein ulex+ (nonzero) pixels in the resulting image multiplied by the squared pixel size. The murine vessel area was defined by the number of mouse CD31+ pixels in the resulting image multiplied by the squared pixel size. Lastly, non-perfused human EC-lined vessel area was defined as the number of human CD31+ pixels in the resulting image multiplied by the squared pixel size. Of note, due to the presence of the PGA mesh in grafts explanted at 2 weeks, which presents autofluorescence in both green and blue channels, the infused fluorescein ulex area was quantified from merged images where the autofluorescence of the PGA results in the cyan color, distinct from infused fluorescein ulex+ cells (green) and nuclei (blue) staining. The number of green colored pixels were quantified by defining a color threshold and creating a binarized image of the green colored pixels only. The perfused vessel area was then defined by the number of non-zero pixels in the resulting image multiplied by the squared pixel size. Results are shown from three independent experiments performed with grafts containing cells isolated from different human donors. In the final experiment, all cell populations were derived from a single donor. Three animals per group were evaluated to observe statistical differences.

### 1.10 Statistical analysis

Results are expressed as mean ± standard deviation. Statistical analysis was performed using one-way ANOVA method with Tukey post hoc comparisons. All analyses were conducted with GraphPad Prism v7.0 and statistical significance was claimed when *p* < 0.05.

## RESULTS

### Isolation and culture of human endothelial cells, pericytes, fibroblasts and keratinocytes under xeno-free conditions

To determine the effects of xeno-free conditions on cell isolation and growth, freshly isolated HECFCs were cultured in xeno-free medium. From visual inspection of the cultures, it was apparent that the xeno-free media formulation supported attachment and growth of ECs from single colonies (Fig. 2A). Serially passaged cells were then used for further functional and phenotypic evaluation. ECs cultured under xeno-free conditions proliferated at a faster rate than cells isolated and cultured in EGM2-Mv media (Figure 2B). Quantitative assessment of barrier formation over time confirmed the similarity of xeno-free medium to EGM2-Mv in supporting EC cell growth and intercellular junction formation, resulting in similar barrier electrical current as assessed by TEER levels (Figure 2C). Transient TNF-α stimulation (10 ng/mL) of confluent xenofree ECs resulted in similar disruption of resistance with subsequent recovery of barrier function to HECFCs cultured in EGM2-Mv media (Figure 2D). Xeno-free cultured ECs demonstrated a stable EC phenotype and expression of key surface and junction markers such as CD31, ZO-1, VE-cadherin and claudin-5, as well the intracellular marker von Willebrand factor (Figure 2E, F). Hematopoietic lineage cell populations (CD45+) were not present in cultures grown under xeno-free culture conditions.

**Figure 2.**
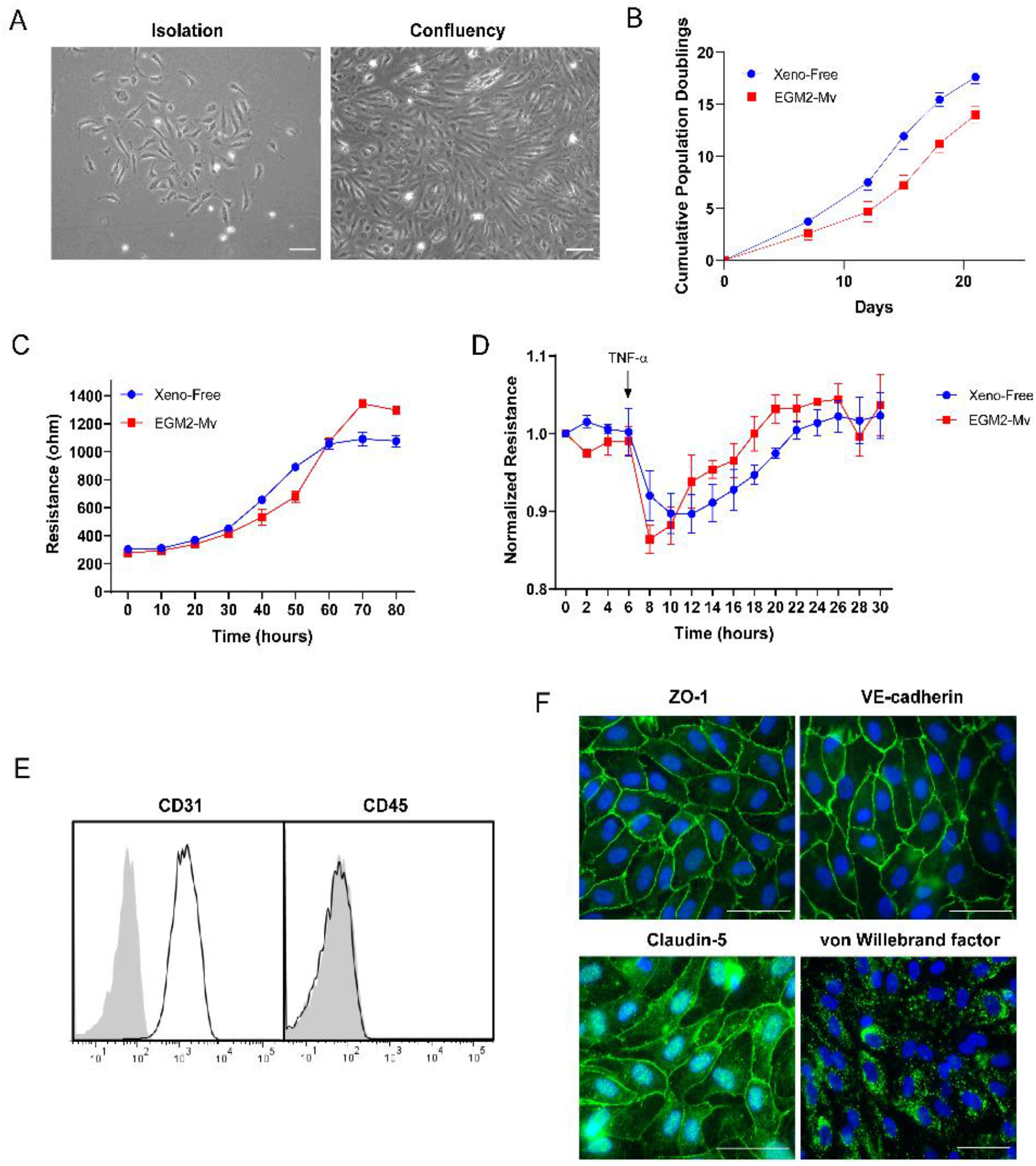
Phenotypic characterization of HECFC-derived ECs cultured under xeno-free conditions compared to cells from the same donor cultured using EGM-2MV medium. **(A)** Live phase-contrast microscopy images of a HECFC-derived EC colony on fibronectin-coated plates after isolation from umbilical cord blood and at confluence. **(B)** The cumulative population doublings of ECs, determined by cell counting, for ECs cultured under xeno-free conditions vs. EGM2-Mv medium. **(C)** Formation of barriers to current flow over time as measured by transendothelial electrical resistance (TEER). **(D)** Barriers are equally disrupted in response to TNF-α (10 ng/mL). **(E)** Surface flow cytometry analysis demonstrating expression of CD31 and absence of CD45 expression. **(F)** Confocal microscopy exhibiting junctional ZO-1, VE-cadherin and claudin-5 staining and intracellular vWF staining in Weibel-Palade bodies. Scale bar: 100 μm. Representative of 3 independent donors.

Human PCs were isolated from placental microvessel fragments by explant outgrowth and serially cultured under xeno-free conditions. Morphological analysis at the time of isolation showed successful attachment of vessel fragments to fibronectin-coated plates (Figure 3A). Xeno-free culture medium showed a faster growth rate and revealed significantly higher cumulative population doublings at the end of passages 2, 3, 4 and 5 compared to PCs cultured in DMEM supplemented with 10% FBS (Figure 3B). The higher cumulative cell number is a result of shorter population doubling time under xeno-free culture conditions. Cultured cells showed positive expression of PDGFR-β, NG2, CD90, and fibroblast activation protein (FAP), but not CD31 (a marker of ECs), CD45 (a marker of hematopoietic cells), or PDGFR-α (a marker of fibroblasts) (Figure 3C). Immunofluorescence microscopy confirmed positive staining of α-smooth muscle actin and FAP (Figure 3D).

**Figure 3.**
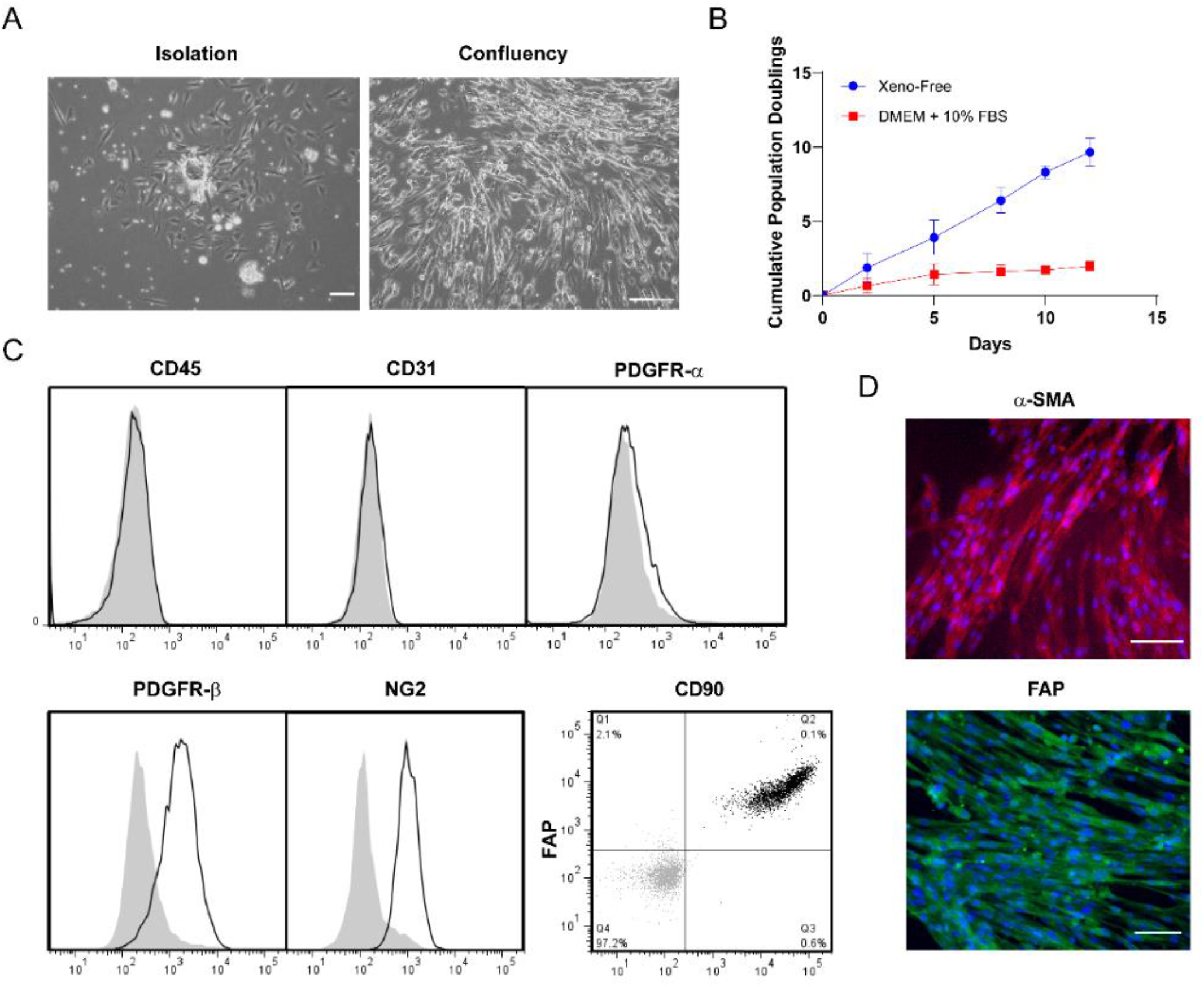
Phenotypic characterization of human placental PCs under xeno-free conditions. **(A)** Live phase-contrast microscopy images of a vessel fragment on a fibronectin-coated plate after isolation from placental tissue and atthe confluency state. **(B)** The cumulative population doublings of PCs cultured under xeno-free conditions is higher than in DMEM supplemented with 10% FBS. **(C)** Flow cytometry analysis of PCs cultured under xeno-free conditions confirmed the expression of PDGFR-α, NG2, CD90 and FAP. Furthermore, these cells lack expression of CD31 and CD45. **(D)** PCs showed positive expression of α-SMA and FAP, as confirmed by immunofluorescence staining. Scale bar: 100 μm. Representative of 3 independent donors.

Human dermal FBs were isolated from foreskin dermis by explant outgrowth and cultured under xeno-free conditions. Cultured cells presented a typical spindle-shaped morphology (Figure 4A) and proliferated at a significantly faster rate than cells isolated and cultured in DMEM supplemented with 10% FBS (Figure 4B).

**Figure 4.**
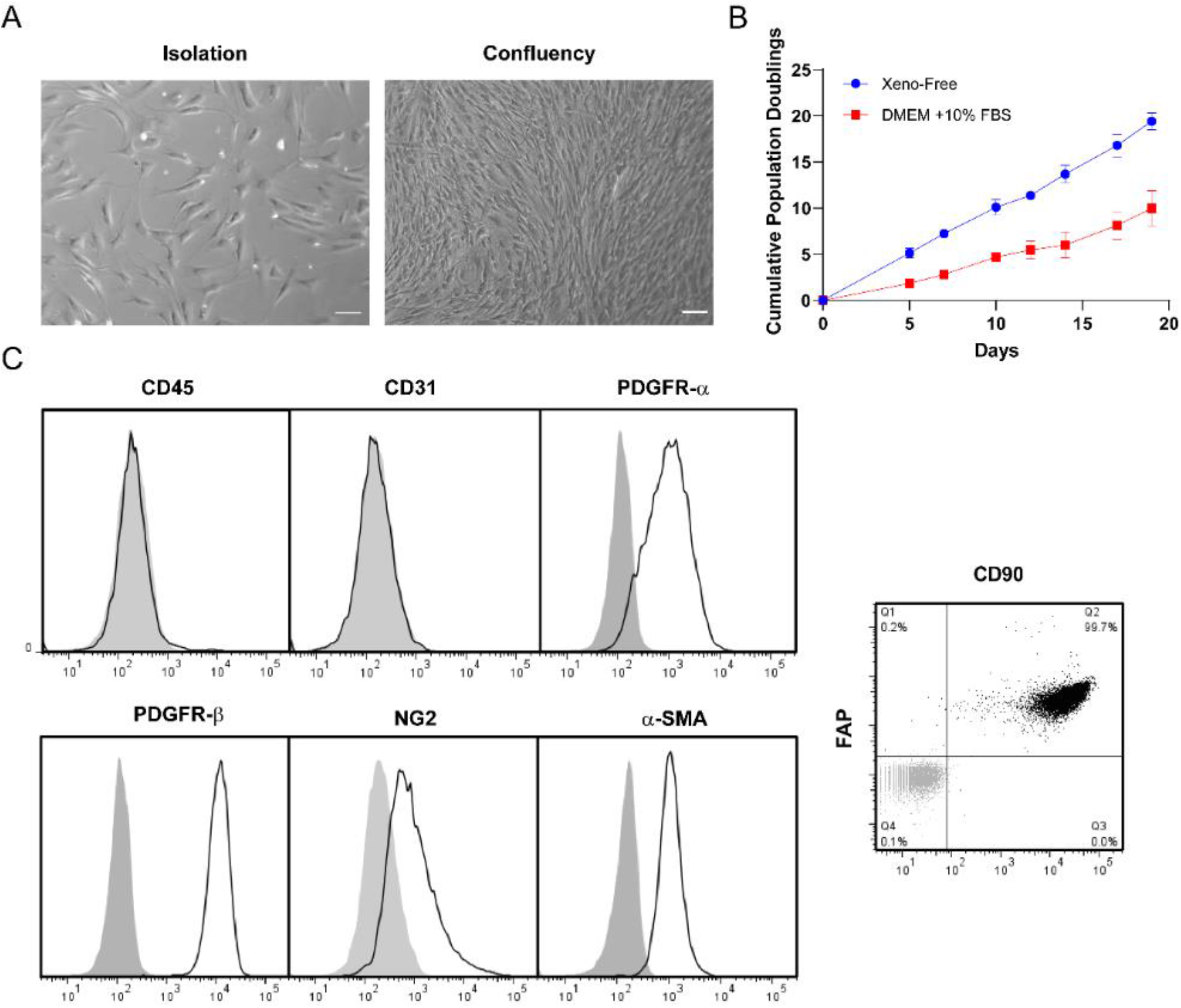
Phenotyping characterization of human dermal FBs under xeno-free conditions. **(A)** Live phase-contrast microscopy images of FBs after isolation from the dermis of donor foreskin and at the confluency state. **(B)** The cumulative population doublings of FBs cultured under xeno-free conditions is higher than in DMEM supplemented with 10% FBS. **(C)** Flow cytometry analysis of FBs cultured under xeno-free conditions confirmed the expression of PDGFR-α, PDGFR-β, NG2, CD90 and FAP. Scale bar: 100 μm. Representative of 3 independent donors.

Fluorescence flow cytometry confirmed that FBs uniformly expressed PDGFR-α, PDGFR-β, FAP, CD90, NG2 and α-SMA, collectively suggesting that cultured dermal FBs isolated under xeno-free conditions display a myofibroblast-like phenotype (Figure 4C) but differ from PCs by expression of PDGFR-α and a more extensively spindled morphology. Neither hematopoietic lineage (CD45+) nor endothelial (CD31+) markers were detected in FBs cultures established and maintained under xeno-free conditions.

Human KCs were isolated from foreskin and cultured under xeno-free conditions. Keratinocytes attached to collagen IV-coated plates presented a typical cobblestone morphology at confluency (Figure 5A).

**Figure 5.**
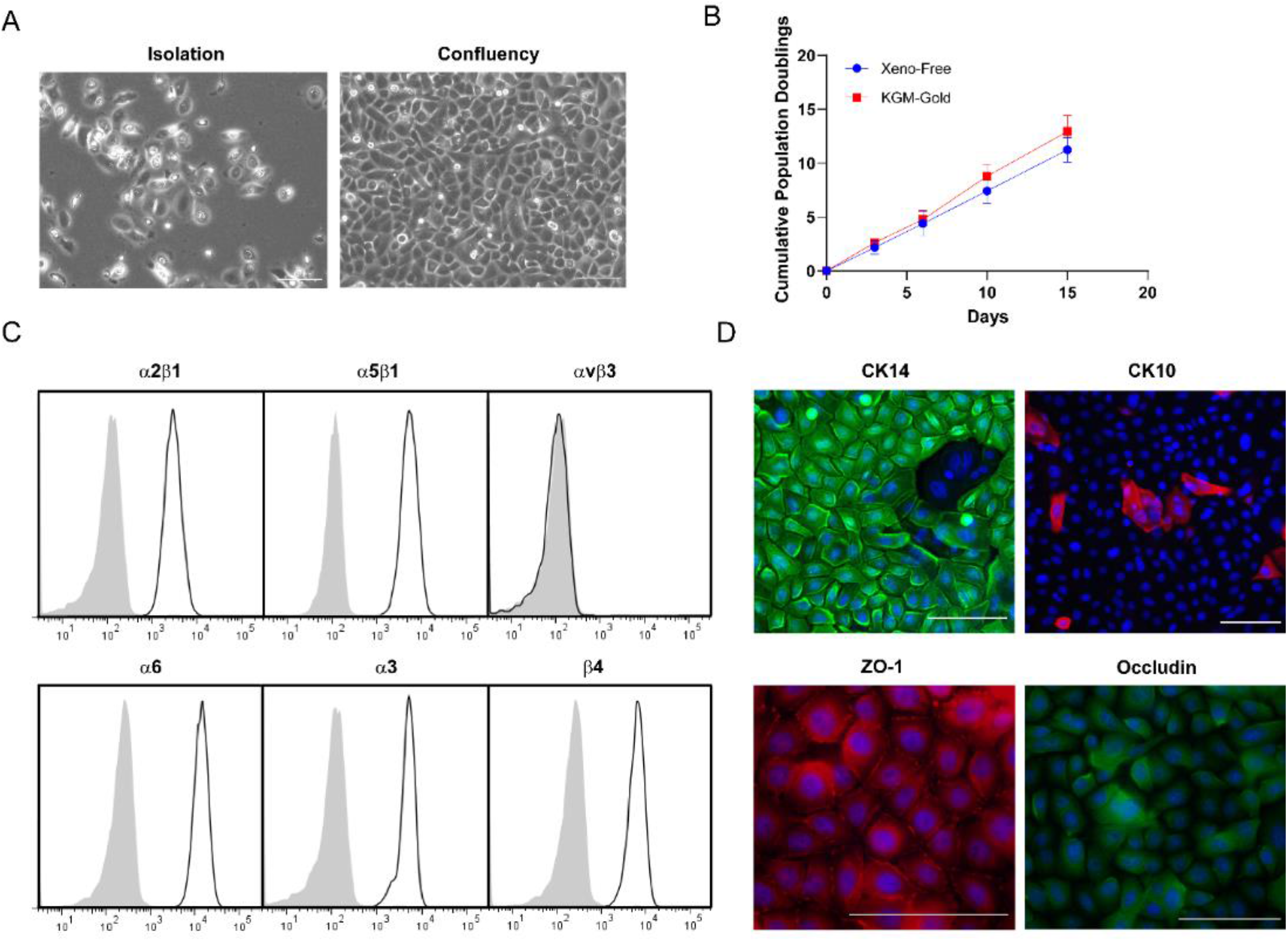
Phenotyping characterization of human keratinocytes under xeno-free conditions. **(A)** Live phase-contrast microscopy images of keratinocytes after isolation from the epidermis of donor foreskin and at the confluency state. **(B)** The cumulative population doublings of keratinocytes cultured under xeno-free conditions is comparable to KGM-Gold medium. **(C)** Flow cytometry analysis confirmed the expression of integrins α2β1, α5β1, α6, α3 and β4, but not αvβ3. **(D)** Confocal microscopy exhibiting CK14, CK10, junctional ZO-1 and intracellular occludin staining. Scale bar: 100 μm. Representative of 3 independent donors.

Quantitative assessment of cell proliferation showed a similar growth rate of keratinocytes cultured in xeno-free medium compared to KGM-Gold medium (Figure 5B). Flow cytometry analysis confirmed the expression of integrins α2β1, α5β1, α6, α3 and β4, but not αvβ3 (absent in normal keratinocytes) (Figure 5C). Moreover, immunofluorescence microscopy showed that most cells at confluency present a basal keratinocyte phenotype (CK14+), while some cells had begun to differentiate into suprabasal keratinocytes (CK10+) (Figure 5D). Keratinocytes monolayers under xeno-free culture conditions express the tight junction protein ZO-1 at the plasma membrane, while occludin expression was found mostly intracellularly.

### Implantation of xeno-free human 3D bioprinted skin grafts onto immunodeficient mice

Bioprinted skin grafts were implanted on the dorsum of immunodeficient mice after 8 days of *in vitro* culture in three independent experiments. In one of these experiments, we used cell populations isolated from a single donor and also generated grafts lacking human ECs and PCs for purposes of assessing the effect of the absence of preformed vascular structures. To assess if human EC-lined vessels are present and become perfused in implanted bioprinted grafts, fluorescent Ulex Europaeus Agglutinin I was injected through the tail vein 30 min before explant. Positive labeling of the ECs assessed in post-harvest tissue sections demonstrated the presence of accessible human EC populations within the dermis of bioprinted grafts indicative of contact with the bloodstream (Figure 6A, lower row). Interestingly, subsequent staining of the sections with human CD31 antibody revealed additional cells, indicating that not all human ECs were present in perfused vessels (Figure 6B). Quantification of infused ulex+ area showed no significant change in the number of perfused human ECs up to 6 weeks post-engraftment (2 weeks: 1.2%; 4 weeks: 1.3%; 6 weeks: 1.8%) (Figure 6C). In contrast, the area of human CD31staining of non-perfused ECs decreased between 2 (1.6%) and 4 (0.74%) weeks but stabilized by 6 weeks (0.77%) (Fig 6D). Skin substitutes explanted 2 weeks post-engraftment showed a high degree of infiltrating mouse vessels, that was more prominent in bioprinted grafts containing human ECs and PCs (14% vs 7.4%) (Figure 6A, S1A). Similarly to non-perfused human vessels, the presence of murine vessels in the dermis of vascularized bioprinted grafts was significantly reduced by 4 weeks (5.5%), but examination of grafts harvested at 6 weeks (4.3%) suggested that mouse vessel number had stabilized (Figure 6E).

**Figure 6.**
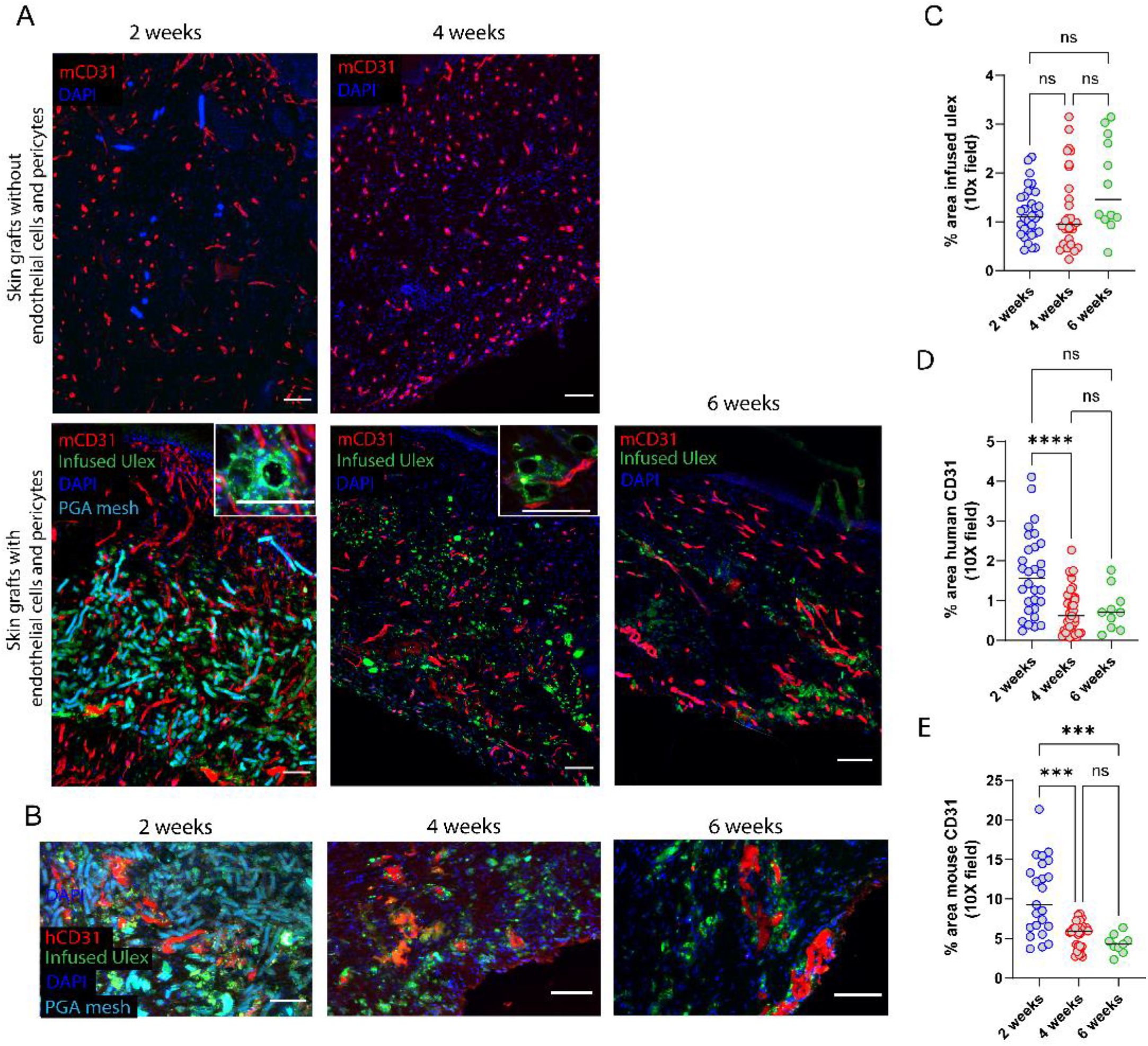
Characterization of graft-derived and host-derived microvessel formation in xeno-free 3D bioprinted skin grafts with and without ECs and PCs 2-, 4- and 6-weeks post-engraftment onto immunodeficient mice. **(A)** Fluorescent mouse CD31 staining depicts higher degree of host angiogenesis (red) and the presence of perfused human EC-lined vessels (infused Ulex stain) in grafts containing ECs and PCs. **(B)** Human CD31 staining shows the presence of non-perfused human ECs (red) surrounded by PGA mesh and perfused human vessels (green). Nuclei are stained blue. Scale bar: 100um. Quantification of area of **(C)** infused ulex, **(D)** human CD31 and **(E)** mouse CD31 at 2-, 4-, and 6-weeks post-implantation. Data are shown for 3 independent experiments. (* indicates p < 0.05, ** indicates p < 0.01, *** indicates p < 0.001, **** indicates p < 0.0001, ns indicates p > 0.05; One-way ANOVA method with Tukey post hoc comparisons).

Previous work both in vitro and in vivo suggests that placental PCs associate with and stabilize microvessels formed by self-assembly of human ECs^19^. Quantification of SMA staining in xeno-grafts 4 weeks post-engraftment showed that about 20% of the total SMA+ cells were associated with vessels in the dermis (Figure S2). This population would include human PCs, but based upon our *in vitro* phenotyping, could also include human FBs, present in ten-fold greater numbers in the bioink, and, since this stain is not human-specific, could also include invading mouse cell populations. Thus it is likely that most, if not all, of the human PCs in the bioink were associated with microvessels.

We anticipated that inclusion of a PGA mesh to impart structural stability to the dermal compartment might elicit a macrophage-rich foreign body type reaction. Indeed, bioprinted grafts with human ECs and PCs showed a high degree of inflammation at 2 weeks following engraftment as shown by the presence of F4/80-expressing mouse macrophages surrounding the PGA mesh (Figure 7, upper row). Four weeks after implantation, the PGA mesh was completely degraded, and grafts were less intensively infiltrated by macrophages, suggesting that the degree of inflammation associated with the presence of the mesh was resolving (Figure 7, lower row). Despite the presence of a macrophage infiltrate, grafts containing human vascular cells did not show evidence of injury. As we have observed in the past ^19^,grafts containing human PCs show a high degree of epidermal organization with keratinocyte maturation and development of rete ridges. Strikingly, bioprinted grafts containing only fibroblasts and PGA mesh in the dermis showed a high degree of hemorrhage, inflammation, and necrosis at 2 weeks (Figure 8, upper row). Histological analysis showed poor epidermal keratinization and presence of necrotic regions surrounding the PGA mesh. At 4 weeks post-engraftment, non-vascularized bioprinted skin grafts showed some degree of recovery and healing as demonstrated by the presence of a scab, re-epithelization (from the wound edges), and a high degree of inflammation compared to grafts containing human ECs and PCs (Figure 8, lower row). The human origin of the healing epidermis was confirmed by involucrin staining, a marker of terminal differentiation of human KCs (Figure S3).

**Figure 7.**
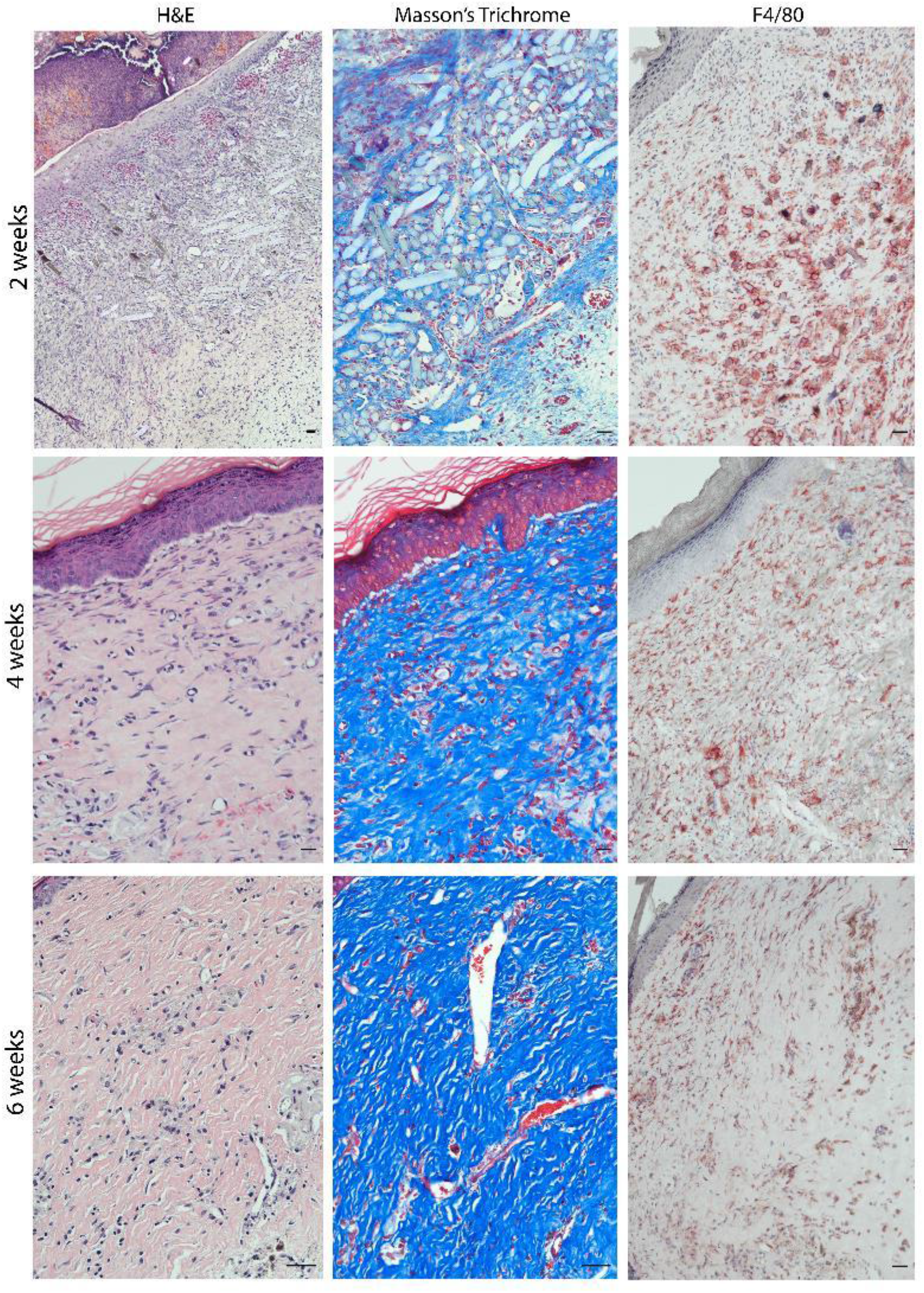
Characterization of xeno-free 3D bioprinted skin grafts with ECs and PCs 2-, 4- and 6-weeks post-engraftment onto immunodeficient mice. Representative images of H&E and Masson’s trichome-stained sections showing degradation of the PGA mesh after 2 weeks. Representative images of F4/80-stained samples (red - mouse macrophages) showing that grafts become less intensively infiltrated overtime. Scale bars: 100 μm

**Figure 8.**
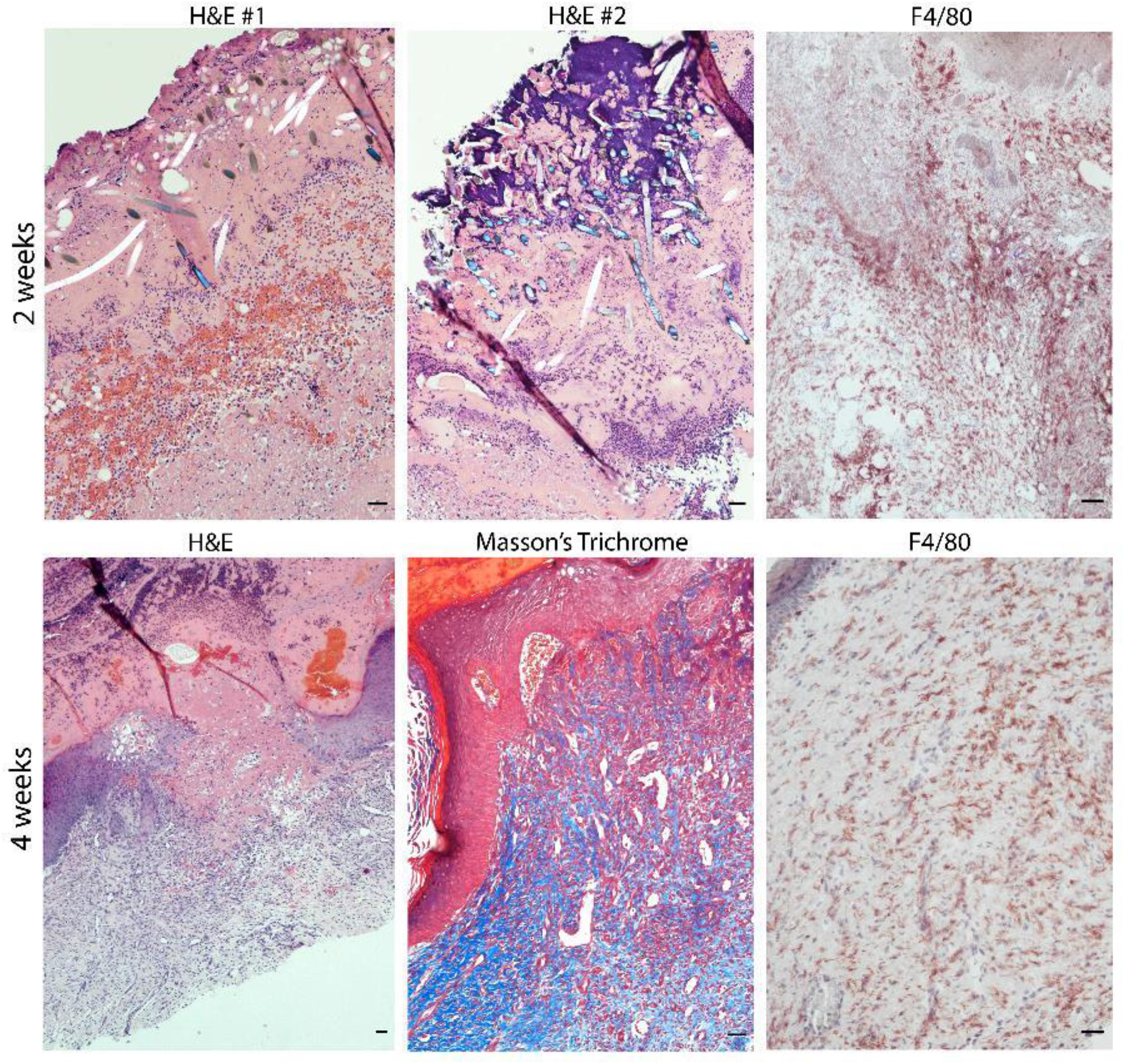
Characterization of xeno-free 3D bioprinted skin grafts without ECs and PCs 2- and 4-weeks post-engraftment onto immunodeficient mice. Representative images of H&E and Masson’s trichome-stained sections showing the presence of PGA mesh and high degree of necrosis in grafts lacking ECs and PCs. Delayed re-epithelization from the wound edges was observed in grafts lacking ECs and PCs 4 weeks post-engraftment. Scale bars: 100 μm

## DISCUSSION

To our knowledge, the creation of a 3D bioprinted skin graft generated under strict xeno-free conditions had not previously been realized. Here, we describe a protocol for successful isolation, culture, and cryopreservation of primary cells, bioink/scaffold development, construct maturation *in vitro*, and successful engraftment on an immunodeficient mouse host. After grafting into a host, the grafts support a vascularized dermis with perfused human EC-lined vessels supported by PCs and a differentiated multi-layered human epidermis. Our results establish the feasibility of bioengineering human tissues and organs under strict xeno-free conditions. This model can also serve as a starting point for introducing additional skin elements, such as melanocytes, skin appendages, and resident leukocytes. This system may also be useful for exploring allogeneic immune responses and their evasion by engrafting a human immune system allogeneic to the skin cell donor.

Our work incorporates several advances in biomaterials. The generation of skin equivalents for transplantation requires the use of multiple isolated cell types and extracellular matrix (ECM) components. The use of collagen type I as a biomaterial has several advantages that include biocompatibility, biodegradability, and the presence of bioactive sites that allow for cell attachment, growth, and differentiation^22^. Currently, most commercially available sources of collagen type I for both research and clinical applications are derived from animal tissues, such as bovine, porcine, rat, and fish^23,24^. Despite some promising clinical outcomes, immunogenic responses (i.e., allergies in 2-4% of the population) and potential for transmission of prion diseases are two major clinical concerns associated with the use of xeno-derived collagen grafts. In this study, we demonstrate that human collagen type I and human fibronectin can be used in the development of a bioink that supports the generation of an implantable and perfusable xeno-free skin graft with a mature stratified epidermis. We also improved the mechanical properties of the 3D printed graft by incorporation of a PGA mesh into the dermal layer. PGA has been available for implantation into humans for over 30 years in the form of surgical sutures^25^. Because it is non-toxic in vivo, this polymer has been widely used as a scaffold for tissue engineering: for example, recent studies utilized PGA as a scaffold for vascular tissue engineering using porcine and human iPSC-derived smooth muscle cells (SMCs)^26,27^. Here, we used a PGA non-woven mesh to provide initial mechanical stability to the graft. The presence of the mesh also allowed for easier handling of the construct during formation and easier suturing of skin grafts at the time of engraftment. Within 4 weeks, the mesh was completely hydrolyzed—in a response mediated by host macrophages—and replaced by new native ECM. The inflammation associated with the presence of the mesh appeared to be largely resolved by 4 and 6 weeks. While there seems to be a high degree of matrix remodeling occurring in the graft at earlier times (2 weeks, for example), no significant changes were observed regarding the number of perfused human vessels that persist in the dermis between four and six weeks. There are some potential disadvantages to using PGA scaffolds in this setting: for example, it has been shown that SMCs grown in high concentrations of PGA breakdown products showed decreased expression of calponin, a marker for SMC differentiation, and decreased SMC proliferation^28^. Although PGA may not be the optimal scaffold for the broad range of vascular tissue engineering applications where complete repopulation of the scaffold with host SMC is desired, it may offer significant advantages for skin tissue engineering applications where graft contraction and scar formation are still current limitations^29–31^.

The major argument for incorporating vascular cells into the dermal layers is to accelerate revascularization of the graft. Grafts produced that lack ECs and PCs underwent extensive early necrosis and keratinocyte loss. Although an epidermis did eventually form in grafts lacking ECs and PCs—likely from surviving human keratinocytes localized near the graft edges—preassembled vessels prevented the early necrotic phase. Furthermore, the presence of ECs and PCs in the bioprinted grafts enhanced host angiogenesis compared to grafts containing only fibroblasts, xeno-free matrix components, and PGA mesh. At 4 weeks-post engraftment about 20% of total α-SMA+ cells associated with vessels. We have previously shown in a co-culture system of RFP-transduced ECs, AmCyan-transduced FBs, and GFP-transduced PCs, that PCs directly associate with ECs while FBs remain randomly distributed^19^. Given the 1:10 ratio of human PCs to FBs in the bioink, 20% of the SMA+ cells number probably reflects the majority of PCs. In addition to vessel stabilization, PCs may have beneficial paracrine effects. Specifically, we and others have previously shown that the presence of PCs induce a higher degree of keratinocyte maturation^19,32,33^.

Other cases of crosstalk between vascular ECs and KCs have been reported^34–36^. It was recently shown that soluble heparin-binding epidermal growth factor (s-HBEGF) released from vascular ECs activates the epidermal growth factor receptor (EGFR) on KCs and promotes stabilization of sirtuin 1 (SIRT1), an essential keratinization inducer^34^. Although the proliferation of human primary KCs with s-HBEGF recombinant protein was almost unchanged in a cell survival assay, a significant increase was detected in the mRNA levels of differentiation markers, corroborating the role of s-HBEGF in facilitating keratinization. These results suggest that that the presence of host and donor ECs in xeno-free skin grafts may not act on KC survival by inducing keratinocyte proliferation, but rather through a paracrine effect on keratinization induction.

Cell sourcing is a critical issue in design of bioinks. Often, the need for a graft is urgent and it must be available “off-the-shelf’ to be most useful. For example, a skin graft could be fabricated using differentiation of patient-derived induced pluripotent stem cells (iPS cells)^37^, but that would require many weeks to accomplish. Consequently, cells used for clinically-relevant graft construction will most likely be derived from another (i.e. allogeneic) individual. Moreover, recent studies have clearly established that human pluripotent stem cell lines display variable capacities to differentiate into specific lineages, even when using iPS cells derived from the same donor, and that the number of cell divisions required creates a mutational burden producing a significant risk for cancer formation^38,39^. Fully differentiated cells from older individuals may senesce. For these reasons, we favor the use of cells from neonates that have reached the stage of committed progenitors or are fully differentiated. Our current results provide proof of principle that this source of cell types is practical for tissue engineering of skin.

Further improvements and modifications to our current methodology are likely to be required for optimization of clinical skin substitutes. Natural human skin is highly immunogenic in patients and in human immune system reconstituted mice models, but avascular synthetic skin grafts, containing only FBs, KCs, and matrix, such as Apligraf™ are not^16^. Our past work has provided a potential mechanism to explain this observation: graft ECs, which largely remain of donor origin in clinical transplantation, become “activated” through deposition of complement and evoke strong allogeneic host immune responses^40–43^. We have proposed that modification of graft ECs to reduce their immunogenicity could reduce the frequency and intensity of rejection without increasing (or possibly even reducing) immunosuppression^44,45^. For example, we found that elimination of MHC class I and class II molecule expression on human ECs, achieved using CRISPR/Cas 9 technology, can diminish recognition of human ECs by both T cells and DSA, and did not cause “missing-self’ recognition by NK cells^46,47^. Furthermore, implantation of self-assembled perfusable networks generated with double knockout (DKO) class I and II MHC - EC into a human immune system mouse model, proved to be sufficient to eliminate rejection. We anticipate that future incorporation of genetically altered cells in bioprinted skin grafts—using cells that are designed to evade alloimmune responses under xeno-free conditions—will allow the generation of an “off-the shelf’ vascularized, allogenic, bioprinted skin graft that can evade rejection or minimize the degree of immunosuppression. Although there is strong evidence that ECs play an important role in triggering human allograft rejection, it is unknown if or which other cell types within a human graft are significant contributors to rejection and thus must also be treated to protect the graft. The experiments with Apligraf cited above appear to argue against an important role for KCs and FBs, but it is equally possible that the lack of vascularization simply kept circulating lymphocytes from contacting these other cells. Interpreting the role of different cell types in bioengineered grafts requires that they come from the same donor and share MHC alleles if the results are to be applied to natural organs. Here, we have developed an implantable bioprinted skin equivalent in which all of the cell populations used were derived from the same neonatal male donor, using cord blood as a source of ECs, placenta as a source of PCs, and foreskin (from neonatal circumcision) as a source of FBs and KCs. The development of this model provides an opportunity to study rejection of bioengineered grafts and to better understand mechanisms of alloimmunity in the context of bioengineered tissues and organs.

While organ transplantation is the single most effective treatment for end stage organ failure, it still requires the use of immunosuppressive regimens with significant side effects, including direct toxicities and increased susceptibility to infections and cancer due to immunological responses of the host to allogeneic tissues^48^. One promising solution is manufacturing of replacement organs through tissue engineering, where significant advances have been made through 3D printing^49–51^. 3D printing offers the opportunity to produce complex macro- and microstructures with different cell types that have been gene-edited or drug-loaded to alter immunogenicity or antigenicity. Reducing rejection will not only enable bioengineered organs to function for longer, but also minimize a need for second transplants. Reduction of the incidence and severity of allograft rejection could both significantly improve current transplant outcomes and increase the likelihood of developing successful therapies to replace multiple tissues and organs.

## ACKNOWLEDGMENTS

This work was supported by NIH grants R01-HL085416 and R21-AI159580. T.B. was supported by the PhRMA Foundation’s Postdoctoral Fellowship in Translational Medicine.

## AVAILABILITY OF DATA AND MATERIALS

All data generated and analyzed in this study are available from the corresponding author on request.

## ETHICS APPROVAL AND CONSENT TO PARTICIPATE

Not applicable.

## CONSENT FOR PUBLICATION

Not applicable.

