## Supplemental figures for "3D bioprinting of an implantable xeno-free vascularized human skin graft"

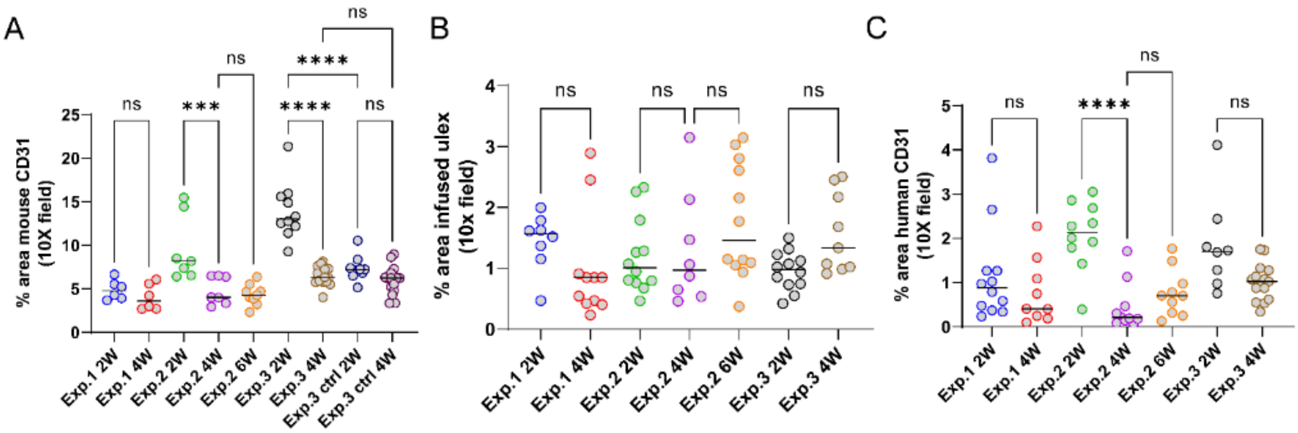


**Figure S1– Quantification of area of (A) mouse CD31, (B) infused ulex and (C) human CD31 at 2-, 4-, and 6-weeks post-implantation.** (* indicates $p < 0.05$, ** indicates $p < 0.01$, *** indicates $p < 0.001$, **** indicates $p < 0.0001$, ns indicates $p > 0.05$; One-way ANOVA method with Tukey post hoc comparisons).

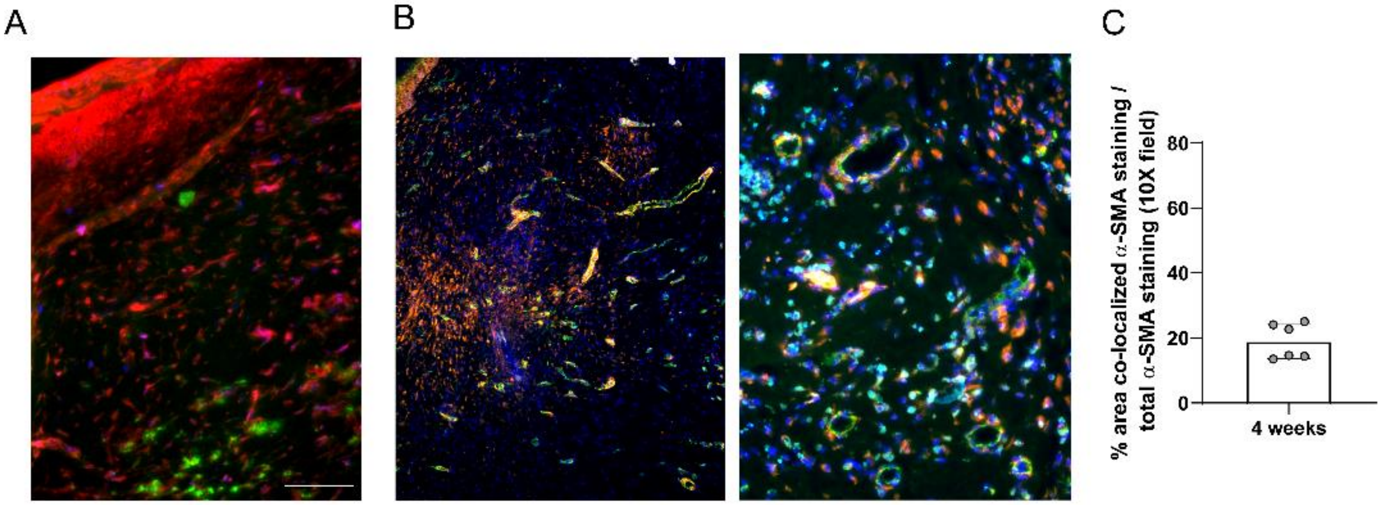


**Figure S2– (A)** Human cells expressing MHC class I are present in the epidermis and dermis of xeno-free grafts 4 weeks post-engraftment. **(B)** Alpha-SMA-positive PCs associate with murine (green) and human ECs (cyan). **(C)** Quantification of PCs associated with vessels in xeno-grafts 4 post-implantation. Scale bars: 100 μm.

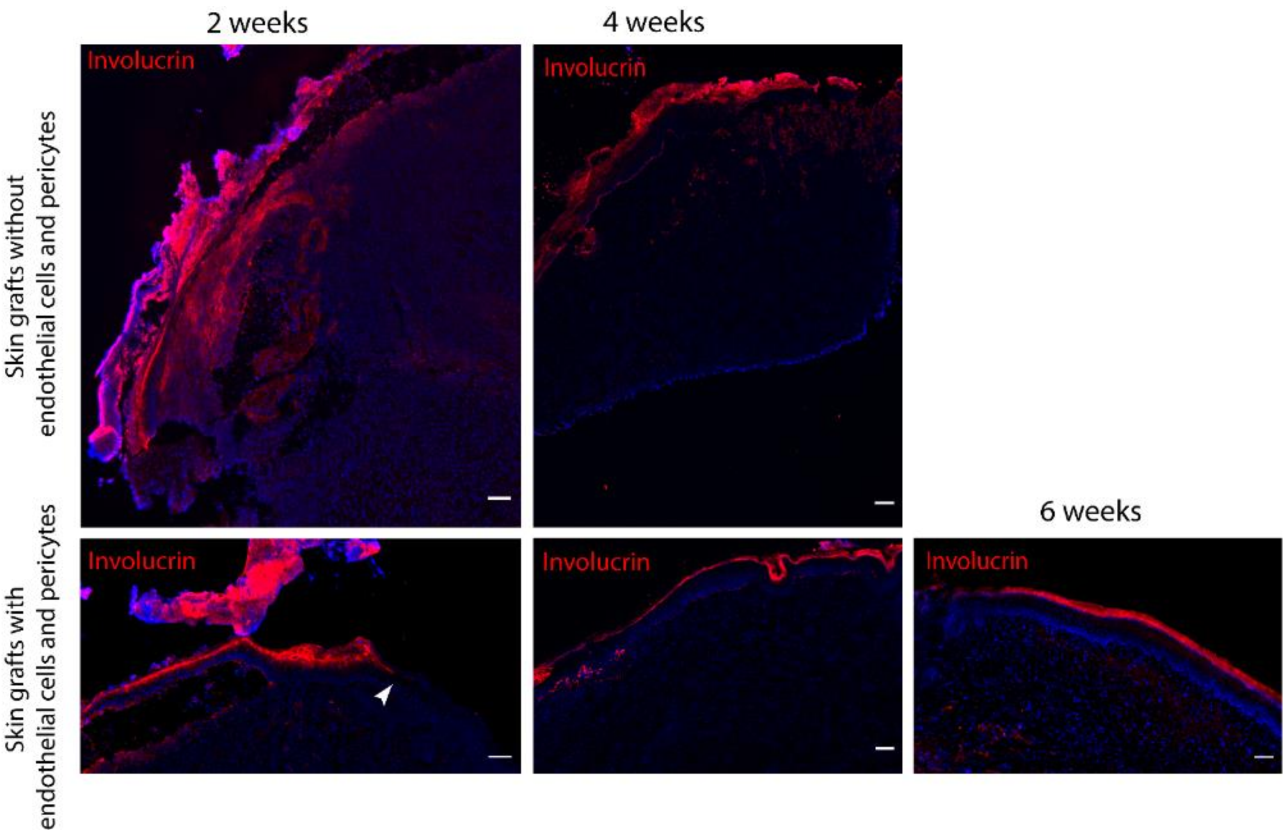


**Figure S3 – Epidermis of bioprinted xeno-free skin grafts with (lower row) and without ECs and PCs (upper row) is of human origin at 2-, 4- and 6-weeks post-engraftment**. Immunofluorescence staining of human involucrin showing that epidermis of 3D bioprinted skin is human. White arrow points to the edge of the wound. Scale bars: 100 um.
